# Histone modification crosstalk between host and pathogen

**DOI:** 10.64898/2026.07.13.738199

**Authors:** Shantinique S. Miller, Joel A. Hrit, Scott B. Rothbart, Evan J. Worden

## Abstract

Bacterial pathogens modulate host cell physiology by secreting effector proteins that rewire host signaling pathways. A subset of these effectors directly modify host chromatin to reprogram gene expression and promote infection. While these enzymes are thought to function autonomously, the extent to which the host epigenetic landscape regulates their activity remains largely unknown. RomA and its homolog LegAs4 are Set domain-containing lysine methyltransferases from *Legionella pneumophila* that methylate histone H3 at lysine 14 (H3K14) to suppress host immune responses and enhance intracellular bacterial replication. Here, we demonstrate that RomA activity is constrained by pre-existing host histone post-translational modifications (PTMs) through multiple layers of histone PTM crosstalk. RomA selectively binds and methylates unmodified histone H3 tails and is inhibited by histone PTMs associated with active transcription, including H3K4 trimethylation, H3K4 acetylation, and H4K12 mono-methylation. We identify both cis- and trans-histone regulatory mechanisms, whereby unmodified H3K4 and H3K14 must reside on the same H3 tail to support RomA activity, while H4K12me1 inhibits RomA across the nucleosome. Notably, cryo-EM analysis and biochemical data reveal that RomA does not engage the nucleosome acidic patch but instead associates flexibly through histone tails. Together, these findings establish the host epigenetic regulation of bacterial effectors as a fundamental and previously unrecognized layer of host-pathogen interactions.

**SIGNIFICANCE:** Bacterial pathogens reprogram host gene expression by delivering effector proteins that modify chromatin, but it is not known how the host epigenetic environment impacts effector function. Here, we show that the Legionella effector RomA senses and responds to the host’s existing epigenetic landscape and is selectively active only in specific chromatin contexts through mechanisms resembling those used by eukaryotic chromatin regulators. Notably, we uncover that RomA utilizes cis-histone and trans-histone crosstalk mechanisms previously observed only in eukaryotic systems. These reveals an unexpected form of host-pathogen crosstalk in which bacterial effector activity can be constrained by host epigenetic modifications.

## INTRODUCTION

Intracellular bacterial pathogens have co-evolved with their hosts over millions of years to acquire a specialized set of effector proteins which allow them to escape immune detection and proliferate^1^. These pathogens use various protein secretion systems to transport effectors into the host cytosol, which then directly interact with host cell machinery to alter eukaryotic signaling pathways^1^. Secreted effectors have been shown to modulate diverse eukaryotic systems ranging from membrane structure^2^ and metabolism^3^ to ubiquitination^4^. Many of these effectors contain domains that are homologous to eukaryotic proteins, likely originating through horizontal gene transfer with their hosts^5^. One class of bacterial effectors, termed nucleomodulins, are specifically targeted to the host cell nucleus to regulate host gene expression^6^. A subset of these nucleomodulins directly bind and modify host chromatin, thereby reprograming host transcription by rewriting the epigenome^7^. Epigenetic nucleomodulins have been identified in various human pathogens including *Bacillus*^*8*^, *Burkholderia*^*9*^, *Chlamydia*^*10*^, *Legionella* ^*6,9*^, and *Mycobacterium*^*11-13*^ indicating that altering the host epigenome is an important and conserved mechanism that has arisen multiple times in pathogenic bacteria. However, despite the importance of this mechanism for pathogen virulence, almost nothing is known about how bacterial effectors interact with the host’s existing epigenome during infection.

In eukaryotes, properly regulated gene expression depends on specific combinations of covalent histone post-translational modifications (PTMs) that form the basis of the signaling network known as the “histone code”^14,15^. In this network, “writer” and “eraser” enzymes attach and remove histone modifications, while “reader” proteins are decoders that bind to specific histone PTMs and either elicit a function directly or recruit other factors^16^. In another layer of complexity, existing histone PTMs can influence the deposition, or recognition, of other histone PTMs in a process termed histone modification crosstalk^17^. RomA and its homolog LegAs4, from *Legionella pneumophila*, are among the most well-studied nucleomodulins and exhibit very tight substrate preferences^6,9^. RomA’s principle enzymatic activity is mono-, di-, and tri-methylation of histone H3 at lysine 14 (H3K14)^6^ . Importantly, RomA’s catalytic activity is required for efficient replication of *L. pneumophila* in human cells^6,9^. RomA-dependent H3K14 methylation is associated with transcriptional downregulation of various immune response genes^6^ and LegAs4 can bind to human heterochromatin protein 1 (HP1) proteins^9^, suggesting that RomA/LegAs4 functions to dull the immune response of the host and thereby help *L. pneumophila* evade detection^6,9^. In un-infected human cells, H3K14 methylation is natively produced at very low levels by the SETD2 and SUV39H1 histone methyltransferases^18^. However, endogenous H3K14 methylation is correlated with increased gene expression and is implicated in DNA repair pathways^18^. It is not clear how RomA-dependent H3K14 methylation exhibits a gene silencing function while endogenous H3K14 methylation correlates with gene activation. One hypothesis is that the increase in H3K14 methylation caused by RomA during *L. pneumophila* infection blocks H3K14 acetylation, a PTM that is highly correlated with active transcription^6^. RomA-dependent methylation of H3K14 may compete with H3K14 acetylation genome-wide^19^, thereby inhibiting transcription by reducing H3K14 acetylation globally. However, another possibility is that H3K14 methylation is specifically deposited at (or excluded from) certain genomic sites due to interactions between RomA and existing histone modifications. In fact, the high density of histone marks in our genome makes it very likely that RomA must interact with nucleosomes that are already modified^20^. However, it is not known if RomA recognizes existing epigenetic signatures or if its activity is modulated by the epigenetic environment of the host cell.

RomA contains an N-terminal catalytic Set domain which is responsible for its H3K14 methylation activity and a C-terminal 3x-Akyrin (Ank) repeat domain (**Fig. 1a**). Ank domains are widely annotated as mediators of protein-protein interactions^21^. Interestingly, the human H3K9 methyltransferases G9A and GLP use Ank domains to “read” existing H3K9 mono-, and di-methyl marks^22,23^. The presence of both Set and Ank domains in RomA suggests that this microbial effector may be able to “read” existing modification states of chromatin as it “writes” H3K14 methylation. If true, this would be the first example of histone modification crosstalk between a bacterial invader and its human host and would greatly expand our understanding of host-pathogen interactions during bacterial infection.

**Figure 1.**
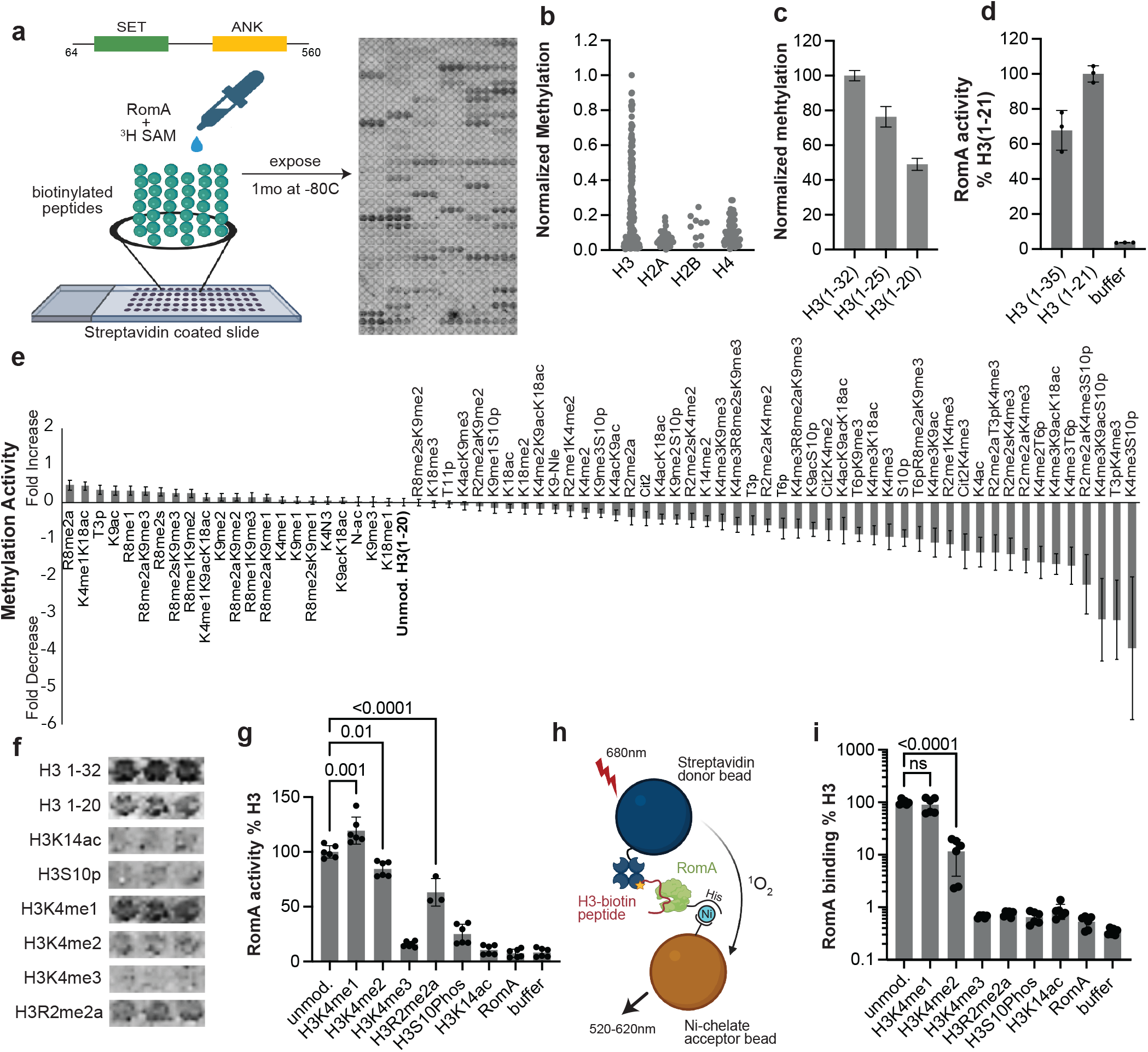
RomA is sensitive to the modification state of histone H3 peptides. a) Domain architecture of the RomA construct used and schematic of the peptide array experiment. b) RomA dependent methylation detected on peptides derived from each histone. c) RomA-dependent H3K14 methylation on unmodified H3 peptides of different lengths. d) MTase-Glo methyltransferase assay on isolated H3 (1-35) and H3 (1-21) peptides. e) Relative methylation of H3(1-21) peptides containing a free H3K14 sidechain and different combinations of modifications. Error bars are standard error of the fold change propagated using taylor expansion f) Raw array methylation data for relevant H3 peptides. g) MTase-glo methyltransferase assays using selected modified H3(1-21) peptide substrates. h) Schematic of Alphascreen binding assays between RomA and biotinylated H3 peptides. i) Relative binding of RomA to selected H3(1-21) peptides. Data shown as mean ±SD. P-values are shown above the indicated samples.

Here, we establish that RomA specifically engages unmodified histone peptides and nucleosomes and that its activity is tightly constrained by the existing epigenetic modifications. RomA is highly sensitive to the modification state of the histone H3 N-terminal tail, with unmodified H3K4me0/me1 serving as preferred substrates, whereas histones containing H3R2 methylation and H3S10 phosphorylation reduce its activity. Importantly, H3K4me3 completely inhibits H3K14 methylation by RomA, revealing tight crosstalk between H3K4me3 and RomA-dependent H3K14me1/me2/me3. RomA reads H3K4 on the same H3 tail as its substrate H3K14 sidechain, showing that RomA is regulated by the H3K4 modification state through a *cis*-histone crosstalk mechanism. Surprisingly, we find that H4K12me1 potently inhibits RomA activity, revealing an additional layer of *trans*-histone crosstalk between H3K14me1/me2/me3 and H4K12me1 that only occurs in the context of intact nucleosomes. This nucleosome dependence, together with the position and identity of the histone tail modifications and the insensitivity of RomA to acidic patch mutations, supports a model in which RomA engages chromatin through histone tails. Our structural data further support this model. Using previously published crystal structures and Alphafold3 (AF3) models of LegAs4 and RomA bound to H3 peptides, we show that RomA binds to the first four amino acids of histone H3, and that H3K4 docks into a pocket on the RomA Ank domain, providing a structural rationale to understand *cis*-histone crosstalk between H3K4me3 and H3K14me1/me2/me3. Together, our findings establish histone modification crosstalk as a fundamental mechanism controlling the activity of epigenetic nucleomodulins and highlight that host chromatin can actively regulates bacterial effector function during infection.

## RESULTS

### RomA catalytic activity is regulated by PTMs on the N-terminus of histone H3

To determine whether RomA catalytic activity is regulated by epigenetic modifications present on host chromatin, we used a modified histone peptide array to measure methylation activity of His-FLAG-RomA(64-560) **(Fig. 1a)** across 276 unique synthetic histone peptides containing individual and combinations of specific PTMs^24^. The RomA(64-560) truncation was chosen as the full-length protein had solubility issues during purification. In this assay, RomA utilizes ^3^H-SAM to deposit ^3^H-methyl onto the immobilized peptides, and methylation is measured using autoradiography (**Fig. 1a**). RomA did not methylate H2A, H2B, or H4 peptides, consistent with the known specificity of RomA for histone H3K14 (**Fig. 1b**)^6^ . Unmodified H3 (amino acids 1-32) showed the highest levels of methylation while shorter H3 peptides (amino acids 1-25 or amino acids 1-20) showed lower methylation levels, indicating longer H3 peptides are better substrates for RomA on the array **(Fig. 1c)**. However, the length dependence of RomA activity disappeared when H3(1-21) and H3(1-35) peptides were methylated in solution (**Fig. 2d**). This indicates that the length dependence of RomA methylation seen in the peptide array is likely due to complex surface interactions, and not due to an underlying preference of RomA for longer H3 peptides. Therefore, to control for any length-dependent effects, we only considered peptides comprising H3(1-20) and normalized our methylation data to the unmodified H3(1-20) peptide. To understand how the H3 modification state impacts RomA activity, we compared methylation levels between unmodified and modified H3 peptides containing an available H3K14 sidechain. RomA methylated distinct H3 peptides to different extents, indicating that RomA is sensitive to the modification state of the H3 tail **(Fig. 1e**). A large subset of H3 modifications inhibited H3K14 methylation by RomA, particularly modifications on H3R2, H3K4, and H3S10. RomA is also sensitive to the specific modification state of H3K4, with RomA activity decreasing in a stepwise manner with each additional methyl group on H3K4 (**Fig. 1e-f**). To orthogonally validate key results from the peptide array, we purchased modified H3(1-21)-biotin peptides and measured RomA activity using the MTase-Glo methyltransferase assay, which measures formation of SAH (**Fig. 1g)**. Peptides containing the H3K4me1 modification stimulated RomA activity relative to an unmodified H3 peptide, while H3K4me2-modified peptides decreased activity. However, peptides containing H3K4me3 reduced RomA activity to ∼5% compared to unmodified H3. In addition, H3 peptides containing R2me2a or S10p reduced RomA activity to ∼50% and 20% compared to the unmodified peptide.

**Figure 2.**
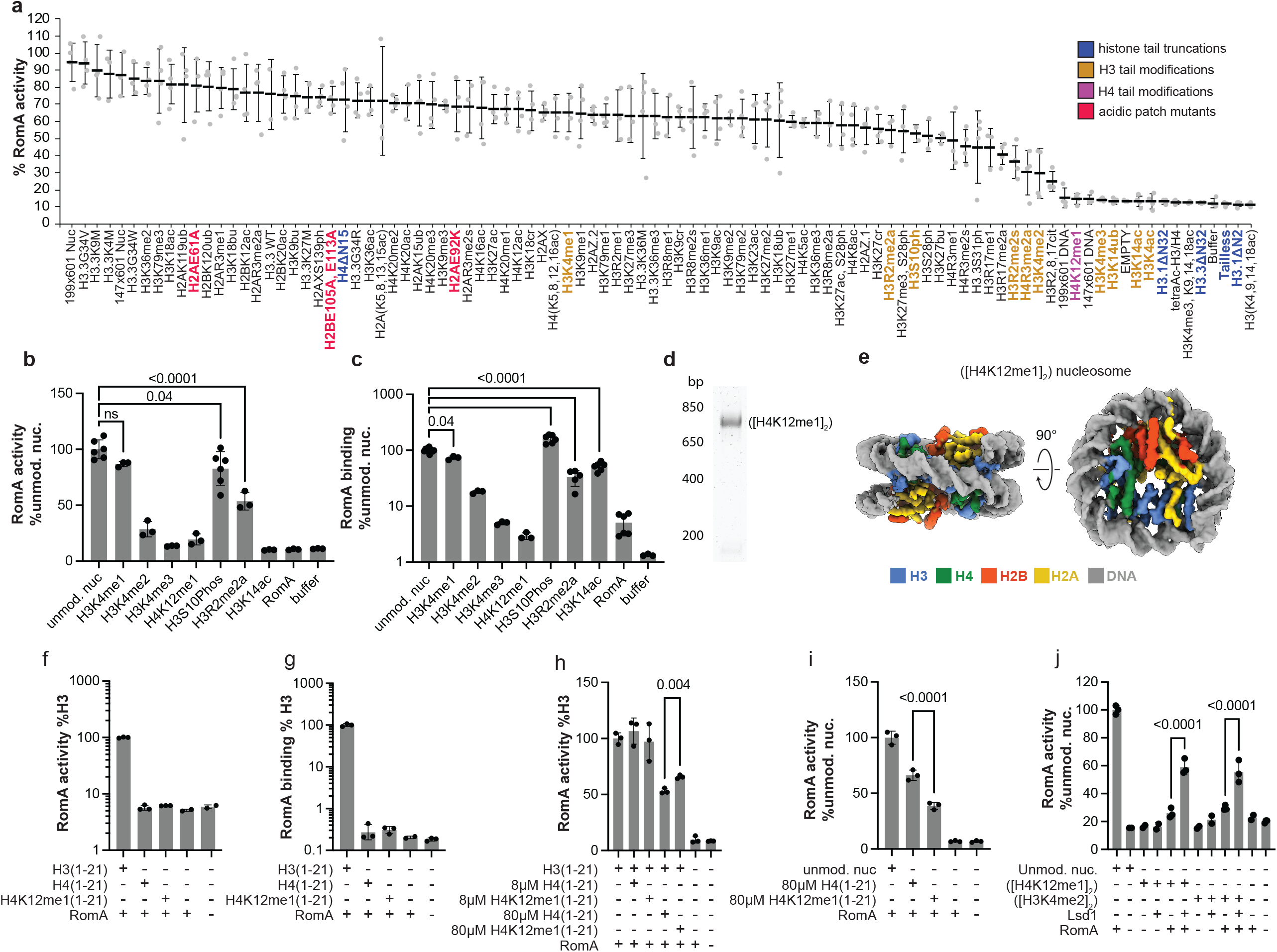
RomA is regulated by histone modifications in the nucleosome context. a) RomA-dependent H3K14 activity on a panel of nucleosome substrates bearing histone variants, mutations, and modifications. Important modifications are colored according to the panel legend. b-c) Normalized MTase-glo activity assays (b) and alpha screen binding assays (c) between His-Flag-RomA and nucleosome substrates bearing single histone modifications. d) Native PAGE gel showing mobility of the H4K12me1 nucleosome. e) Cryo-EM structure of the (H4K12me1]2) nucleosome colored according to the panel legend. f-g) MTase-glo RomA activity assay (f) and alpha screen binding assay (g) using with the indicated H3 and H4 peptide substrates. h) MTase-glo RomA methylation assay on unmodified H3(1-21) in the presence of H4(1-21) peptides. i) MTase-glo RomA activity assay on unmodified nucleosomes in the presence of 80µM H4(1-21) peptides. j) MTase-Glo RomA activity on unmodified, ([H4K12me1]2) and ([H3K4me2]2) nucleosomes before and after treatment with 1µM Lsd1. Data shown as mean ± SD. P-values are shown above the indicated samples.

We next used the C-terminal biotin on the H3 peptides and the N-terminal His affinity tag on His-FLAG-RomA (64-560) to conduct Alpha Screen binding assays (**Fig. 1h-i**). These assays showed that H3K4me2/me3, H3R2me2a and H3S10p modifications reduced the ability of RomA to bind its H3 tail substrate, suggesting that the reduced methylation activity of RomA on these peptides is caused by binding defects. Interestingly, RomA did not bind to an H3K14ac peptide, indicating that interactions with the lysine sidechain are also important for H3 binding. Taken together, these data show that the modification state of the H3 tail, particularly at H3K4, regulates RomA-dependent H3K14 methylation and H3 binding.

To understand if the nucleosome structure is important for RomA activity, we characterized RomA-dependent H3K14 methylation activity using the Captify™ nucleosome panel (EpiCypher), which contains ninety-three homotypic recombinant nucleosomes bearing distinct histone modifications and mutations (**Fig. 2a**). Hereafter, we refer to modified nucleosomes using the standardized nomenclature^25^.Unmodified nucleosomes were the preferred substrates for RomA with all other modified nucleosomes having reduced RomA activity (**Fig. 2a**). In agreement with our peptide array data, the nucleosome panel showed that RomA is particularly sensitive to modifications on the H3 N-terminus. Nucleosomes bearing H3K4me1/me2/me3 modifications showed a stepwise decrease in H3K14 methylation, ([H3K4me1]_2_) > ([H3K4me2]_2_) > ([H3K4me3]_2_), with RomA activity progressively reduced with each additional H3K4 methyl group and completely inhibited on H3K4me3 nucleosomes. ([H3S10p]_2_) and ([H3R2me2a]_2_) reduced RomA activity while ([H3K4ac]_2_), ([H3K14ub]_2_) and N-terminal H3 truncations ([H3.1ΔN2]_2_), ([H3.3ΔN32]_2_), ([H3.1ΔN32]_2_)) completely blocked H3K14 methylation by RomA (**Fig. 2a**). MTase-glo and alpha screen assays, using newly purchased biotinylated nucleosomes (Epicypher), showed that the decreased RomA activity for each modified nucleosome was correlated with reduced RomA binding affinity, except for ([H3S10p]_2_), which had enhanced affinity for RomA (**Fig. 2b-c**). In particular, ([H3K4me3]_2_) completely blocked RomA binding and activity, suggesting that H3K4me3 disrupts an important interface between RomA and the nucleosome. Together, these results show that RomA is highly sensitive to the modification state of the H3 N-terminus, and that RomA distinguishes between different nucleosomes through crosstalk with histone modifications.

### Trans-histone crosstalk between H3K14 methylation and H4K12me1

([H4K12me1]_2_) was among the worst substrates for RomA in the nucleosome panel (**Fig. 2a**). This was unexpected as no interactions between RomA and H4 have been reported and a full deletion of the H4 tail ([H4ΔN15]_2_) only moderately reduced RomA activity (**Fig. 2a**). Notably, H4K12ac did not significantly inhibit RomA activity (**Fig. 2a**) despite occurring on the same sidechain, showing that RomA specifically sensitive to the H4K12me1 modification, not any modification on that residue. To confirm the inhibitory effect of H4K12me1, we purchased fresh ([H4K12me1]_2_) nucleosomes (EpiCypher) and conducted H3K14 methylation and RomA binding assays (**Fig. 2b-c**). In agreement with the results from our nucleosome panel, the H4K12me1 modification potently inhibited RomA activity and completely ablated RomA binding as measured by AlphaScreen. To confirm that the loss of RomA activity is not due to a nucleosome assembly defect, we conducted an electromobility shift assay (EMSA) which showed that the ([H4K12me1]_2_) nucleosome runs at its expected mobility (∼ 700bp, **Fig. 2d**). In addition, we determined a Cryo-EM structure of the un-crosslinked ([H4K12me1]_2_) nucleosome, which showed clear density for all eight histones and fully wrapped DNA, indicating that the nucleosome was assembled correctly (**Fig. 2e, S1, table 1**). MTase-glo and Alpha Screen assays showed that RomA does not methylate or bind H4(1-21) or H4K12me1(1-21) peptides, ruling out a direct association between RomA and the H4 tail in isolation (**Fig. 2f-g**). In addition, an 8-fold molar excess of H4(1-21) or H4K12me1(1-21) peptides (1µM) did not affect RomA activity toward H3(1-21) peptides when added to the methylation reaction while a 640-fold molar excess of the same H4 peptides (80µM) reduced RomA activity by ∼50%, likely due to non-specific effects of the high H4 peptide concentration (**Fig. 2h**). However 80µM of the H4K12me1(1-21) peptide reduced RomA activity on unmodified nucleosomes more than the unmodified H4(1-21) peptide (**Fig 2i**) indicating that the inhibitory effect of H4K12me1 requires the nucleosome. Finally, to understand if removing the H4K12me1 and H4K4me2 modifications could rescue RomA activity, we pre-treated the ([H4K12me1]_2_) and ([H3K4me2]_2_) nucleosomes with a the H3K4-specific lysine demethylase Lsd1 (**Fig. 2j**). Lsd1 has not been shown to demethylate H4K12me, therefore we used a 10-fold molar excess of Lsd1 (1 µM) to favor demethylation of H3K4 and H4K12. Pre-treatment of the ([H4K12me1]_2_) and ([H3K4me2]_2_) nucleosomes with Lsd1 increased RomA activity to ∼60% of the unmodified nucleosome (**Fig. 2j**). We suspect that the incomplete rescue after Lsd1 treatment is due to the high concentration of Lsd1 in the reaction, which may compete with RomA for H3 tail or nucleosome binding. Together, our results show that the inhibitory effect of H4K12me1 is exclusively due to the context of this mark within the nucleosome through a *trans*-histone modification crosstalk mechanism

### Cis-histone crosstalk between H3K14 methylation and H3K4me3

Our biochemical studies clearly show that RomA activity and substrate binding requires unmodified H3K4 and H3K14 residues **(Fig 2b)**. However, the nucleosome contains two copies of histone H3 and it is not clear if these two unmodified residues must be present on the same histone tail to stimulate RomA activity. To address this question, we asked if a nucleosome containing unmodified H3K4 on one tail could stimulate H3K14 methylation on the opposite tail. To test this, we purchased heterotypic ([H3K14ac]·[H3K4me3]) nucleosomes (EpiCypher) in which one H3 tail is modified with H3K14ac and the other is modified with H3K4me3 **(Fig 3a)**. In this substrate, neither H3 tail contains unmodified H3K4 and unmodified H3K14, so RomA can only methylate H3K14 if it recognizes H3K4 in the opposite H3 tail. RomA efficiently methylated unmodified nucleosomes, but did not methylate([H3K14ac]_2_), ([H3K4me3]_2_) or ([H3K14ac]·[H3K4me3]) nucleosomes **(Fig 3b)**. Together, these data indicate that RomA reads the modification state of H3K4 and methylates H3K14 on the same H3 tail through cis-histone modification crosstalk.

**Figure 3:**
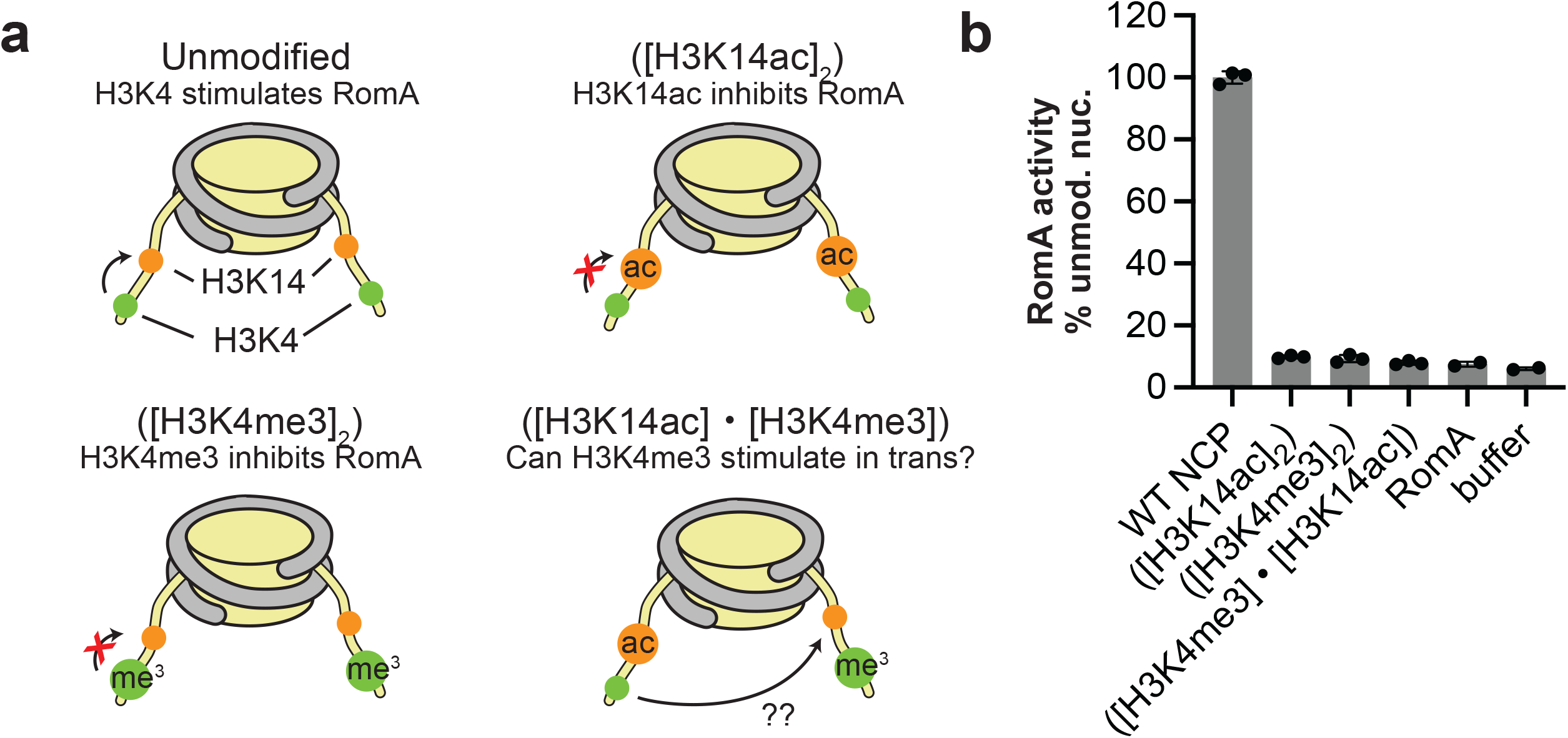
RomA recognizes H3K4 and methylates H3K14 in cis. a) Schematic diagrams of nucleosome substrates used to evaluate cis- vs. trans-histone crosstalk between H3K4 and H3K14 methylation by RomA. b) RomA activity on the nucleosome substrates in a. Data shown as mean ±SD

### RomA does not bind the nucleosome acidic patch

To understand how RomA interacts with the nucleosome, we determined a Cryo-EM structure full-length RomA, fused to an N-terminal MBP solubility tag, bound to a mono-nucleosome where H3K14 was mutated to the non-native amino acid norleucine ([H3K4Nle]_2_) (**Fig. 4, S2-3, table 1**). In the presence of the SAM cofactor, norleucine mutations become high-affinity inhibitors of lysine methyltransferases that trap the enzymes in a substrate-bound state^26,27^. The resulting Cryo-EM maps show clear density for the nucleosome with an extra, diffuse density on top of the nucleosome (**Fig. 4a**). This density is the approximate size of the folded part of LegAS4 (**Fig. 4a, S4a**), so we interpret this diffuse density to be RomA. However, the RomA density could not be clearly resolved even with extensive Cryo-EM processing (**Fig. S3**) or when using multiple distinct RomA/nucleosome preparations, suggesting that RomA does not associate with the folded nucleosome core in a defined conformation. Interestingly, we consistently observed that DNA unwinding from the nucleosome coincided with appearance of the RomA density and that nucleosomes with fully wrapped DNA did not show RomA density, suggesting that RomA binding to the nucleosome core may require structural rearrangement of the DNA (**Fig. S4b-c**). In addition, local refinement of the nucleosome core to high resolution did not show any extra density near the acidic patch residues of H2A and H2B that could be attributed to RomA (**Fig. 4b,c**). This suggests that RomA does not interact with the nucleosome acidic patch, a conserved interaction surface used by most nucleosome-interacting proteins^28,29^. In support of this, RomA activity was not meaningfully impaired on ([H2AE61A]2) or ([H2BE105A]_2_, [E113A]_2_) nucleosomes containing acidic patch mutations (**Fig. 2a**). We validated these observations by purchasing ([H2AE92K]_2_), ([H2AE61A]_2_), and ([H2BE105A]_2_, [E113A]_2_) nucleosomes (EpiCypher) containing acidic patch point mutations (**Fig. 4d-e**). RomA binding and activity was not affected by any acidic patch mutations, indicating that RomA uses a distinct binding interface that is independent of the acidic patch to interact with nucleosomes. Therefore, our biochemical and structural data suggest that RomA flexibly associates with the nucleosome using the histone tails without making stable interactions with the nucleosome core.

**Figure 4:**
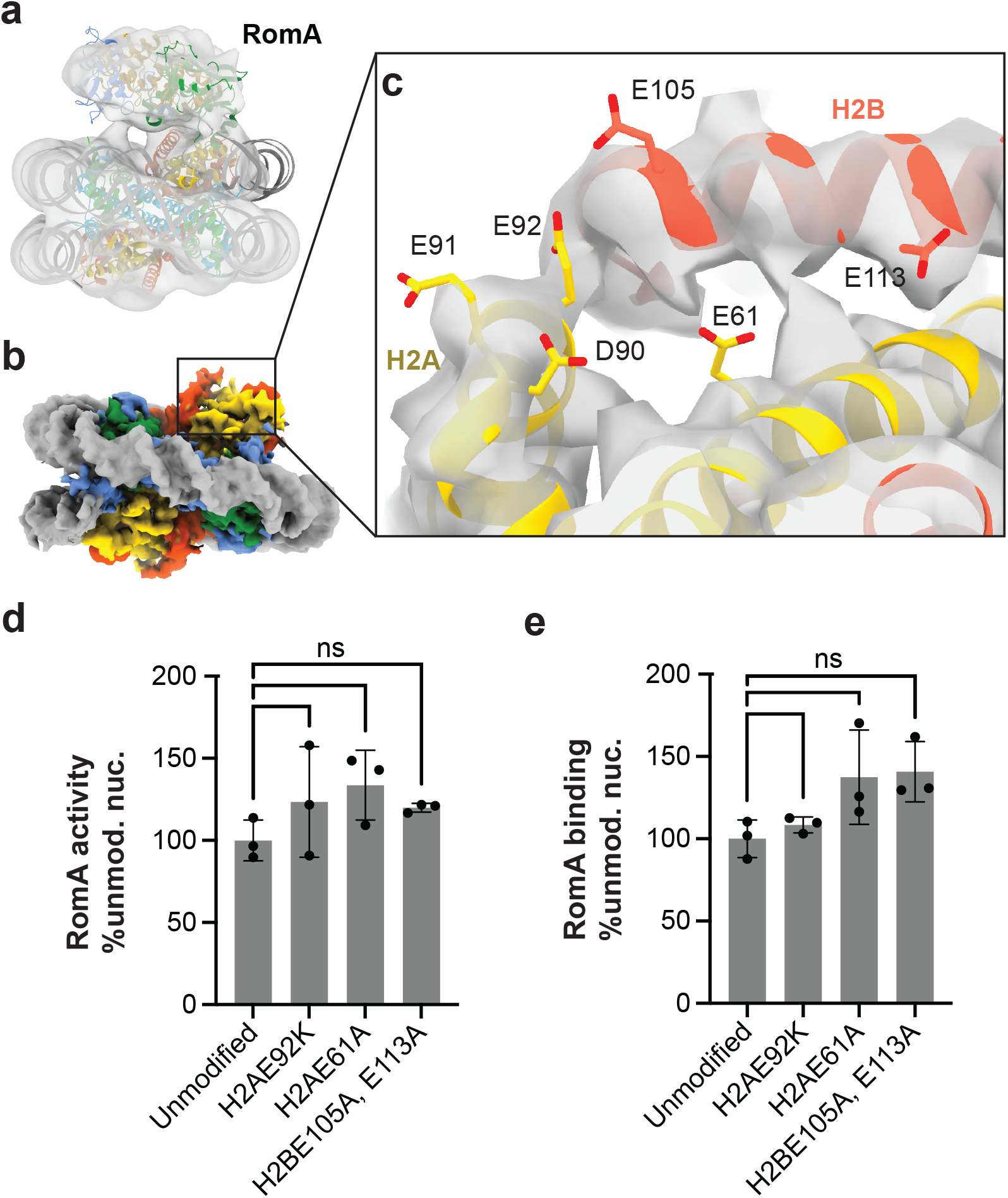
RomA does not bind to the acidic patch. a) Cryo-EM map of the RomA-nucleosome complex, low pass filtered to 15A for clarity and displayed at a contour level 0.0618. Atomic models of LegAS4 (PDB: 5CZY) and the nucleosome (PDB: 8BYJ) are docked into the cryo-EM map. b) Focused refinement map of the nucleosome in the RomA-nucleosome complex and displayed at a contour of 0.198. The map is colored according to Fig 2e. c) Close up view of nucleosome acidic patch in the RomA nucleosome complex. Acidic patch residues are shown as sticks and the map is shown as a semi-transparent gray surface. The crystal structure of a nucleosome wrapped by 601 DNA (PDB ID: 8CZE) is docked into the cryo-EM map. d-e) MTase-glo RomA activity assay (d) and alpha screen binding assay (e) on WT and acidic patch mutated nucleosomes. Data shown as mean ±SD. P-values are shown above the indicated samples.

### Structural basis for crosstalk between H3K4 and H3K14 methylation

No structures of RomA are currently available, but structures of the closely related homolog LegAs4 (96% sequence identity) have been determined in isolation and bound to H3 (1-12) or H3 (3-17) peptides^30^. However, the H3 (1-12) and H3 (3-17) peptides are not substrates for RomA or LegAs4 because they lack either an intact H3 N-terminus or unmodified H3K14 sidechain. Therefore, we generated a model of LegAs4 bound to an H3 (1-17) peptide by superimposing structures of LegAs4 bound to the H3 (1-12) or H3 (3-17) peptides. This composite model shows that the H3 N-terminus docks between the Set, linker and Ank domains, H3K4 inserts into a pocket in the Ank domain and H3K14 inserts into the LegAs4 active site (**Fig. 5a-b**). Alphafold3 predictions of the RomA H3(1-21) complex show a highly similar binding configuration (**Fig. S5a-b**) and suggest that the composite model closely approximates a structure of RomA/LegAs4 bound to an H3 substrate peptide. Histone H3(7-9) could not be modeled confidently, and the Alphafold3 PLDDT score for residues in this region are low (**Fig. S5c**), indicating that the N- and C-terminal portions of the H3 tail may be flexibly linked when bound to RomA/LegAs4. Because the N- and C-terminal portions of the H3 tail may not be rigidly connected when bound to LegAs4, we asked if RomA can methylate H3(11-21) when H3(1-10) is provided in trans. MTase-glo activity assays show that RomA cannot methylate H3K14 in H3(11-21) alone or in the presence of H3(1-10) **(Fig 5c)**. Therefore, RomA requires an intact, continuous, unmodified H3 tail for methylation activity.

**Figure 5:**
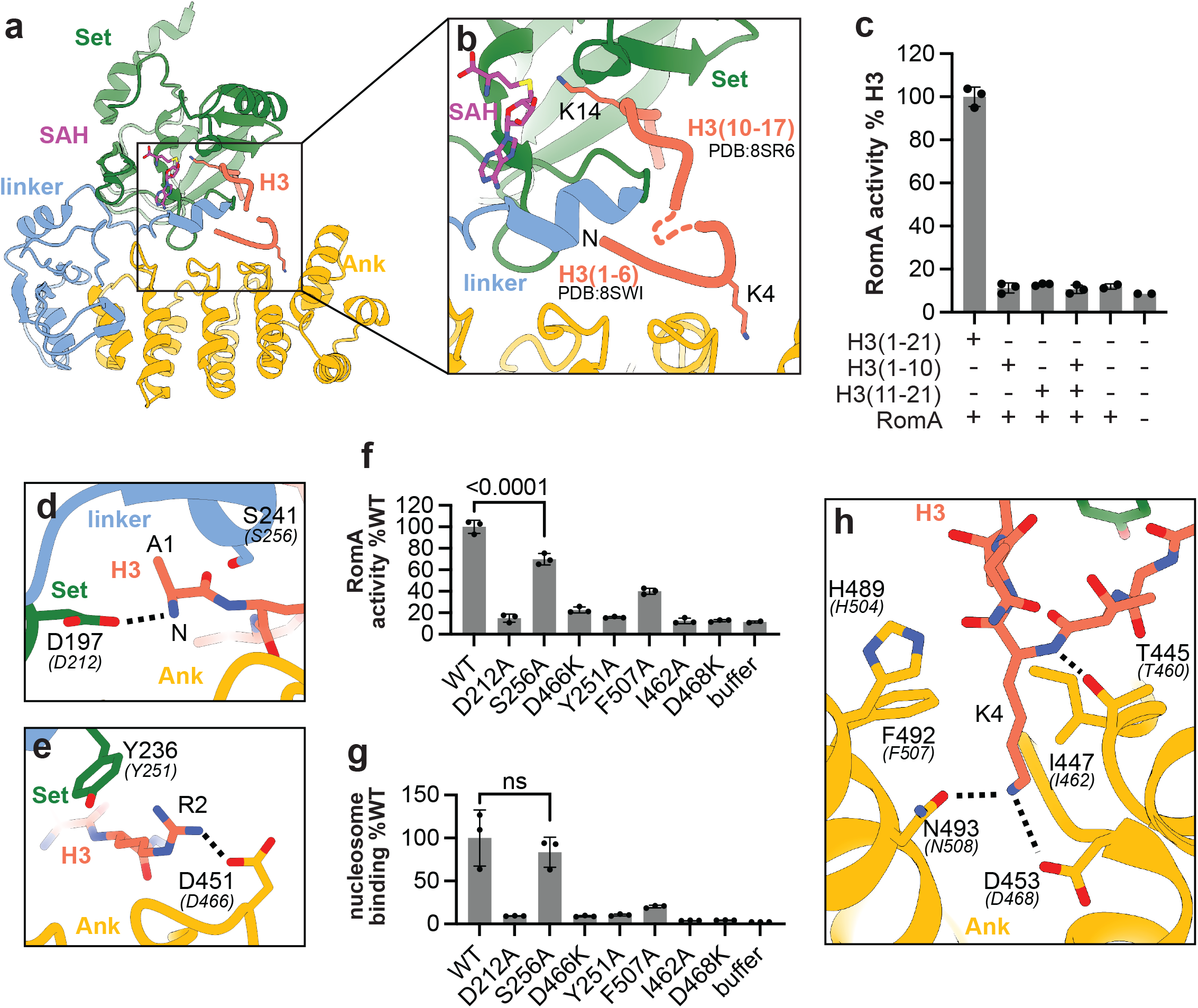
Structural basis for crosstalk between H3K4 and H3K14me1-3 by RomA/LegAs4. a) Superimposition of LegAs4 bound to H3(1-12, PDB:8SWI) and LegAs4 bound to H3(3-17, PDB: 8SR6). LegAs4 from PDB 8SWI is shown in ribbon representation, H3(1-6, PDB:8SWI) and H3(10-17, PDB 8SR6) are shown as red coil. b) close up view of the LegAs4 H3 binding site showing the composite H3 peptide from PDBs 8SWI and 8SR6. H3K4 and H3K14 are shown in stick representation. H3(7-9) are shown as a dashed line. c) MTase-Glo RomA activity assays on a truncated H3(1-10), H3(11-21) peptides. d) Close up view of the interaction between LegAs4 and the H3 N-terminus. Corresponding residues in RomA are shown in the parentheses. e) Close up view of the interaction between LegAs4 and H3R2. f) MTase-Glo activity assays using unmodified nucleosomes and the indicated RomA mutants. g) Alpha screen binding assays between unmodified nucleosomes and the indicated RomA mutants. h) Close up view of the H3K4 binding pocket. Interacting residues are shown in stick representation, important contacts are indicated with dashed black lines. Data shown as mean ±SD. P-values are shown above the indicated samples.

Contacts between the H3 N-terminus and RomA are critical for RomA activity and explain how PTMs on H3 affect H3K14 methylation. The H3 N-terminus interacts with the LegAs4 D197 (RomA D212) (**Fig. 5d**, corresponding RomA residues are shown in parentheses) and is close to LegAs4 S241 (S256). In addition, LegAs4 D451 (D466) forms a salt bridge with H3R2 while LegAs4 Y236 (Y251) sandwiches H3R2 between the Set and Ank domains (**Fig. 5e**). The RomA (D212A) mutation abolished H3K14 methylation and nucleosome binding while RomA(S256A) reduced methylation without affecting binding, showing that contacts with RomA D212 are critical for coordinating the H3 N-terminus. In addition, RomA(Y251A) and RomA(D466K) mutants completely blocked H3K14 methylation and nucleosome binding (**Fig. 5f-g**). This is consistent with reduced H3K14 methylation on H3R2me2a nucleosomes (**Fig. 2b**), as these methyl marks could block interactions between H3R2 and RomA. H3K4 docks into a pocket in the Ank domain (**Fig. 5h**) lined by F492 and I447 (F507 and I462) which contact the aliphatic portion of H4K4 while D453 and N493 (D468 and N508) reside at the bottom of the H3K4 binding pocket and coordinate the H3K4 lysine sidechain (**Fig. 5h**). The F507A mutation reduced RomA activity and binding, while the I462A and D468A mutations completely abolished it (**Fig. 5f-g**). Therefore, these residues are critical for H3K4 recognition, and it follows that H3K4 methylation disrupts these interaction to reduce RomA activity. Indeed the AF3 prediction of a complex between RomA and an H3K4me3 peptide shows that H3K4 no longer docks onto the ANK domain (**Fig. S5d**). A BioRxiv preprint posted during revision of this manuscript reaches many of the same conclusions about the structural basis for crosstalk between RomA and H3 tail modifications^31^. Together, the AF3 predictions and prior crystal structures of LegAs4 bound to non-substrate H3 peptides^30^, explain how RomA reads out the modification state of the H3 tail to regulate its H3K14 methylation activity.

## Discussion

Our work establishes that RomA-dependent H3K14 methylation is regulated by the existing epigenetic modifications through cis- and trans-histone crosstalk mechanisms. To methylate H3K14, RomA directly interrogates the modification state of the H3 and H4 tails, which are located on opposite sides of the nucleosome. Therefore, the crosstalk interactions we describe here enable RomA to read out the epigenetic status of the nucleosome as a whole. This allows RomA to achieve exquisite control of its catalytic activity and ensures that H3K14 methylation is only deposited on *bona fide* substrate nucleosomes. The exceptionally tight control of RomA activity mirrors that of eukaryotic chromatin writers that are natively regulated by cis-, or trans-histone modification crosstalk mechanisms. For example, Polycomb Repressive Complex 2 (PRC2) is allosterically stimulated by H3K27me3 from a different H3 tail (trans-histone crosstalk)^32,33^, while H3K4me3 and H3K36me3 inhibit PRC2 activity when present in the same H3 tail as the H3K27 substrate sidechain (cis-histone crosstalk)^34^ . RomA conditioning its activity to the specific modification state of the nucleosome likely ensures that H3K14 methylation is correctly deposited within host chromatin during infection.

While many bacterial effectors hijack host signaling and metabolic pathways^11,35,36^, RomA represents the first example of a bacterial effector that directly reads the modification state of host chromatin during infection. Chromatinized DNA and post-translational epigenetic modifications are evolutionary hallmarks of eukaryotic cells that are generally absent in bacteria. Therefore, it was surprising that RomA is so strongly regulated by histone modifications. Thus, RomA is the product of a remarkable host-pathogen adaptation where *L. pneumophila* evolved to utilize epigenetic pathways of its hosts that do not exist in its native cellular context. Multiple epigenetic nucleomodulins have been identified across bacterial pathogens^7,8,10,11,35-37^ and many may be similarly sensitive to the epigenetic contexts of chromatin. Therefore, histone modification crosstalk is a previously unrecognized strategy by which bacteria may exploit the host epigenetic landscape to guide reprogramming during infection.

RomA appears to be particularly sensitive to histone modifications associated with active transcription. H3K4me3 is the primary hallmark of active promoters in eukaryotic cells^38-40^ and H4K12me1 is preferentially enriched at active promoters and has been associated with active transcription^41,42^. Similarly, H4R3me2a correlates with active chromatin and promotes recruitment of transcriptional coactivators such as CBP/p300^43,44^. RomA’s sensitivity to H3K4me3, H4K12me1 and H4R3me2a indicates that its activity is anticorrelated with active chromatin states and may instead be most active at regions that do not have an active chromatin signature. During infection, RomA localizes to transcription start sites of immune response genes and promotes the spread of H3K14me early during infection^6^. Interestingly, the close RomA homolog, LegAS4 binds to HP1, a central mediator of heterochromatin formation and transcriptional silencing. This indicates that RomA may function in concert with repressive chromatin pathways^9^. Therefore a function of, RomA may be to silence immune genes before their activation in response to infection. In this context, RomA-dependent H3K14 methylation may function in parallel with HP1-associated mechanisms to reinforce repressive chromatin architecture at specific immune gene promoters.

While the mechanistic basis for RomA inhibition by H3 tail modifications is readily apparent, the molecular mechanism for RomA inhibition by H4K12me1 modifications is much less clear. The effect of H4K12me1 on RomA is likely indirect because RomA is only inhibited in the context of the nucleosome, does not require the H4 tail for activity (**Fig. 2a**), is not inhibited by other modifications on H4K12 (**Fig. 2a**) and does not bind or methylate H4 tail peptides (**Fig. 2f-g**). We speculate that the H4K12me1 modification may alter dynamic properties of the nucleosome such as DNA flexibility or histone H3 tail accessibility to inhibit RomA activity.

Such indirect, histone modification crosstalk effects are well-documented and have been previously described for eukaryotic epigenetic writer complexes. For example, H3K36me3 inhibits PRC2 indirectly by altering contacts between the H3 tail and DNA. This small change causes PCR2 to disengage from the H3 tail and bind nucleosomes dynamically^45^. In addition, H2BK34ub stimulates Dot1L indirectly by propagating small conformational changes through the nucleosome to the H3K79 reside, biasing it to an active state conformation^46^. Therefore we expect that H4K12me1 may occlude the H3 tail from RomA or alter the dynamics of DNA wrapping, which may be important for binding as the RomA density only appears on nucleosomes with altered DNA wrapping (**Fig. S4**).

Together, our findings show how bacterial effector proteins exploit host histone modifications to regulate their own activity. Rather than indiscriminately methylating host chromatin, RomA operates as a highly sensitive writer whose activity is tightly constrained by the existing epigenetic modifications. By integrating histone modification crosstalk at multiple sites, RomA evaluates the epigenetic context before catalysis. This multilayered recognition likely ensures that H3K14 methylation occurs within narrowly defined epigenetic contexts, potentially enabling selective repression of immune genes during infection while avoiding widespread disruption of transcription. Collectively, these features position RomA as an evolutionary specialized effector that exploits chromatin not just as its substrate but as a regulator of its own enzymatic activity.

## Author Contributions

SSM and JAH conducted experiments, SSM collected and processed Cryo-EM data. SBR and EJW advised the research, SSM and EJW conceived the study, SSM wrote the manuscript with input from EJW, SBR and JAH.

## Acknowledgements

SBR and EJW acknowledge support from the National Institutes of Health R35GM152184 (SBR) and 5R21AI173758 (EJW). We acknowledge Cryo-EM Core (RRID:SCR_023210), especially Xing Meng, for their assistance with data collection.

## Methods

### Cloning and Protein Expression

RomA (64–560) and corresponding point mutants were cloned into the pET-28A expression vector, fused to an N-terminal 6xHis-FLAG tandem affinity tag. *E. coli* Rosetta2(DE3) pLysS or BL21(DE3) codon plus cells were transformed with the RomA constructs for recombinant protein expression. For expression of RomA (64-560), 2L of LB supplemented with kanamycin and chloramphenicol was inoculated with *E* .*coli* transformed with the RomA expression vector and grown at 37 °C with shaking until OD_600_ = 0.6–0.8, at which point protein expression was induced by the addition of 1 mM isopropyl β-D-1-thiogalactopyranoside (IPTG). Following induction, the culture was incubated overnight at 18 °C. For expression of RomA mutants 200mL of Autoinduction System 1 media (Millipore) supplemented with kanamycin and chloramphenicol was inoculated with E. coli bearing a mutant RomA expression vector and grown overnight at 37 °C followed by 5-6hr at 18 °C. Cells were harvested by centrifugation at 4,000 rpm for 20 minutes. The cell pellets were resuspended in lysis buffer containing 500 mM NaCl, 30 mM HEPES pH 7.5, 20 mM imidazole, 5% glycerol, and 1 mM β-mercaptoethanol (BME). Resuspended pellets were flash-frozen and stored at −80 °C until further processing.

Full-length RomA(1–560), was cloned into a custom golden gate donor vector based on pET21 with an N-terminal maltose binding protein (MBP) tag. *E. coli* BL21(DE3) pRARE cells were transformed with the RomA constructs for recombinant protein expression. For expression of MBP-RomA(1-560), 2L of LB supplemented with ampicillin and chloramphenicol was inoculated with E.coli transformed with the RomA expression vector and grown at 37 °C with shaking until OD_600_ = 0.6–0.8, at which point protein expression was induced by the addition of 1 mM isopropyl β-D-1-thiogalactopyranoside (IPTG). Following induction, the culture was incubated overnight at 18 °C. The cell pellets were resuspended in lysis buffer containing 500 mM NaCl, 30 mM HEPES pH 7.5, 5% glycerol, and 1 mM β-mercaptoethanol (BME). Resuspended pellets were flash-frozen and stored at −80 °C until further processing.

### Protein Purification

For WT His-FLAG RomA (64-560), cell pellets were thawed at 25 °C and cooled on ice. Once thawed, lysis buffer (500 mM NaCl, 30 mM HEPES pH 7.5, 20 mM imidazole, 5% glycerol, and 1 mM β-mercaptoethanol) was added to a final volume of 150 mL and supplemented with 20 mg of lysozyme. Cells were lysed by sonication on ice for a total time of 12 minutes using cycles of 15 seconds on and 45 seconds off at 45% amplitude. The lysate was clarified by centrifugation at 17,000 rpm for 30 minutes at 4 °C. The clarified lysate was applied to a 5 mL HisTrap HP column (Cytiva) pre-equilibrated with lysis buffer. Affinity purification was performed at 4 °C. Bound proteins were eluted using a linear gradient of lysis buffer containing 500mM imidazole. Fractions containing His-FLAG RomA (64-560) were pooled and concentrated using a 30 kDa molecular weight cutoff spin concentrator (Millipore). The concentrated protein sample was separated by size-exclusion chromatography using a Superdex 200 pg 16/600 HiLoad column (Cytiva) equilibrated in SEC buffer (500 mM NaCl, 30 mM HEPES pH 7.5, 5% glycerol, and 1 mM DTT). Peak fractions were collected, concentrated as needed, flash-frozen, and stored at −80 °C until further use.

For MBP-RomA (1-560), cell pellets were thawed at 25 °C and cooled on ice. Once thawed, lysis buffer A (500 mM NaCl, 30 mM HEPES pH 7.5, 5% glycerol, and 1 mM β-mercaptoethanol) was added to a final volume of 150 mL and supplemented with 20 mg of lysozyme. Cells were lysed by sonication on ice for a total time of 12 minutes using cycles of 15 seconds on and 45 seconds off at 45% amplitude. The lysate was clarified by centrifugation at 17,000 rpm for 30 minutes at 4 °C. The clarified lysate was applied to a 5 mL MBP Trap column (Cytiva) pre-equilibrated with lysis buffer A. Affinity purification was performed at 4 °C. Bound proteins were eluted using lysis buffer A containing 20 mM maltose. Fractions containing MBP-RomA (1-560) were identified, pooled, and concentrated using a 30 kDa molecular weight cutoff spin concentrator (Millipore). The concentrated protein sample was separated by size-exclusion chromatography using a Superdex 200 pg 16/600 HiLoad column (Cytiva) equilibrated in SEC buffer (500 mM NaCl, 30 mM HEPES pH 7.5, 5% glycerol, and 1 mM DTT). Peak fractions were collected, concentrated as needed, flash-frozen, and stored at −80 °C until further use.

For His-FLAG RomA (64-560) point mutants, cell pellets were thawed at 25 °C and cooled on ice. Once thawed lysis buffer was added to a final volume of 10mL. Cells were lysed by sonication on ice for a total time of 3 minutes using cycles of 5 seconds and 15 seconds off at 30% amplitude. The lysate was clarified by centrifugation at 4500 rpm for 30 minutes at 4 °C. The clarified lysate was applied to 200uL of Ni-NTA beads pre-equilibrated with lysis buffer and incubated for 1 hour at 4 °C. After the binding was complete, the beads were washed with lysis buffer. Bound proteins were then eluted with lysis buffer containing 500mM imidazole and spin concentrated to the desired concentration.

### Mtase-Glo Methyltransferase Assays

Methyltransferase activity assays were performed using the MTase-Glo™ Methyltransferase Assay (Promega), a bioluminescence-based assay that quantitatively measures the formation of S-adenosyl-L-homocysteine (SAH), a product of SAM-dependent methylation reactions. All methyltransferase assays were conducted using His-FLAG RomA (64-560) and its corresponding mutants. All reactions were carried out in reaction buffer containing 50 mM NaCl, 30 mM HEPES (pH 7.5), 1 mM dithiothreitol (DTT), and 0.2 mg/mL bovine serum albumin (BSA).

For peptide-methylation assays, reactions contained 250 nM histone peptide substrate, 10 nM RomA, and 20 µM S-adenosyl-L-methionine (SAM). For nucleosome-methylation assays, 125nM nucleosome core particles (Epicypher) were used with 10 nM RomA and 20 µM SAM, except for the acidic patch mutants where 250nM nucleosome was used. For the assays that involved analyzing RomA activity on H3 Peptides in the presence of H4 and H4K12me1 peptides, reactions contained 10 nM RomA, 250 nM H3 peptide, 2000nM H4 or H4K12me1 peptide and 20 uM SAM. All reactions were assembled to a final volume of 12 µL and incubated at room temperature for 30 minutes.

For methylation of nucleosomes pre-treated with His-LSD1 (151-852, Reaction Biology), 1uM of LSD1 was pre-incubated with 100nM nucleosome at room temperature for 4 hours in a final volume of 20uL. After incubation, 20uL of 20nM RomA and 40uM SAM was added to the reaction and incubated for an additional 30 mins at room temperature, yielding a final volume of 40uL.

Following incubation, 10 µL of each reaction was transferred to an OptiPlate™ 384-well plate, and 2 µL of 6X MTase-Glo™ Reagent was added to each well. Plates were centrifuged at 1,000 rpm for 2 minutes, mixed for 2 minutes, and incubated for an additional 30 minutes at room temperature. Subsequently, 12 µL of MTase-Glo™ Detection Solution was added to each well, followed by centrifugation, mixing, and a final 30-minute incubation. Luminescence was measured using a NEO2 multimode plate reader (BioTek).

### Methylation Assay on Modified Peptide Arrays

The histone peptide array was constructed as previously described^47^. Slides were equilibrated by incubation in 5 mL of reaction buffer for 10 minutes at room temperature. Methyltransferase reactions were prepared in a total volume of 200 µL per condition. Reactions contained His-FLAG RomA (64-560) at a final concentration of 100 nM, supplemented with 5 µCi of tritiated S-adenosyl-L-methionine (^3^H-SAM). A control reaction containing 5 µCi ^3^H-SAM in the absence of enzyme was included to assess background signal. Following equilibration, each reaction mixture was pipetted onto an individual peptide array slide and evenly distributed by covering with a glass coverslip.

Slides were incubated overnight at room temperature in a humidified chamber to prevent evaporation. The following day, slides were washed five times for 5 minutes each with 1X phosphate-buffered saline (PBS) to remove unincorporated ^3^H-SAM. Slides were dried by centrifugation at 800 × g for 1 minute. Dried slides were placed into transparent sheet protectors, secured in an autoradiography cassette, and exposed to X-ray film at −80 °C for one month. After exposure, the sealed cassette was equilibrated to room temperature prior to film development.

### Alpha Screen Assays

Alpha Screen binding assays were performed using the Alpha Screen® assay kit (Revvity) containing streptavidin-coated donor beads and nickel chelate (Ni-NTA) acceptor beads. In this assay, His-FLAG RomA (64-560) and the biotinylated nucleosomes were used to enable proximity-based signal detection. All reactions were assembled in Alpha Screen assay buffer consisting of 25 mM HEPES (pH 7.5), 250 mM NaCl, and 0.05% NP-40. His-FLAG RomA (64-560) (100 nM) was incubated with biotinylated substrate (30 nM) in a total volume of 10 µL in an Alpha Plate™ for 1 hour at room temperature to allow complex formation. Following incubation, 5uL of streptavidin donor beads and Ni–NTA acceptor beads were added to each well to a final bead concentration of 20 µg/mL. All bead handling and incubations were performed under low-light conditions to prevent photo-bleaching, in accordance with manufacturer’s recommendations. The final reactions were incubated for an additional 30 minutes at room temperature in the dark. Alpha Screen signals were measured using a NEO2 multimode plate reader (BioTek) equipped with AlphaScreen-compatible excitation and emission filter settings. Signal intensity was reported as AlphaScreen counts.

### Statistical analysis

Results from the modified peptide array are plotted as fold change and the error bars correspond to the standard error of the mean, propagated from the standard error of the raw data using Taylor expansion. All other binding and activity assays are shown as the mean ± a standard deviation of the data. P-values were calculated using the one-way ANOVA test in prism.

### Cloning and Expression of H3K14Nle Histone

To generate histone H3 containing the unnatural amino acid norleucine (Nle) at position K14, histone H3 was cloned into the pQE-81L vector and lysine 14 was mutated to methionine (K14M). For protein expression, the methionine auxotrophic strain E. coli B834(DE3)pLysS was transformed with the pQE-81L(H3K14M) plasmid and grown at 37 °C in M9 minimal medium supplemented with all amino acids. Cells were cultured to an OD_600_ of 0.6, harvested by centrifugation, and washed three times with M9 medium lacking methionine as has been described previously^26^. The cell pellet was resuspended in 25 mL of fresh methionine-free M9 medium supplemented with either 20 mg of norleucine. Protein expression was induced with 1 mM IPTG, and cultures were incubated at 37 °C for 2 h. Cells were then harvested by centrifugation, and the H3K14Nle histone was purified as previously described by Luger et al^48.^

### Nucleosome Reconstitution

Nucleosome reconstitution methods were adapted from Luger et al^48^ . In brief, histone octamers were assembled by combining individually unfolded histone proteins and refolding them together by dialysis into refolding buffer (10mM Tris pH7.5, 2M NaCl, 1mM EDTA and 5mM BME). Octamers were purified by size-exclusion chromatography using a HiLoad Superdex 200 16/600 pg column (Cytiva) and fractions histone octamers were pooled, concentrated and stored at 4 °C until use. Nucleosomes were reconstituted by combining histone octamers with purified Widom 601 DNA in RB-high (10mM Tris pH7.5, 2M KCl, 1mM EDTA and 1mM DTT), followed by gradient dialysis over 18h into RB-low (10mM Tris pH7.5, 0.25M KCl, 1mM EDTA and 1mM DTT). Fully assembled nucleosomes were captured using a TSK DEAE-5PW HPLC column in TES-250 buffer (10mM Tris pH7.5, 0.25M KCl, 0.5mM EDTA and 1mM DTT) and eluted with a gradient of TES-600 buffer (10mM Tris pH7.5, 0.6M KCl, 0.5mM EDTA and 1mM DTT). Pure nucleosome fractions were concentrated and stored at 4 °C until use.

### EM Sample Prep

EM sample preparation utilized MBP-RomA (1–560). RomA was incubated with H3K14Nle-containing nucleosomes at a 10:1 molar ratio (10uM RomA and 1uM nucleosomes) in dilution buffer containing 50 mM NaCl, 30 mM HEPES (pH 7.5), and 1 mM dithiothreitol (DTT), supplemented with 350 uM (SAM). Complexes were incubated for 40 min at 4 °C to allow adequate binding. Following incubation, ∼300 µL of the RomA– nucleosome complex was pipetted onto a 10–30% glycerol gradient prepared in buffer containing 30 mM HEPES (pH 7.5) and 50 mM NaCl and supplemented with 0.07% glutaraldehyde. Gradients were subjected to ultracentrifugation using a SW41 rotor at 32,000 rpm for 18 hours at 4 °C, as described previously^49^. The sample tubes were fractionated from bottom to top, and aliquots of each fraction were analyzed by 7% native polyacrylamide gel electrophoresis (PAGE) to assess complex formation. Fractions corresponding to the fully assembled RomA–nucleosome complex were identified, pooled, and concentrated to ∼0.3mg/mL for subsequent Cryo-EM analysis. Grids used for analysis were Quantifoil R 1.2/1.3 gold grids.

### Data Collection

Cryo-EM data for the complex between RomA and the H3K14NLE nucleosome were collected at the Van Andel Institute Cryo-Electron Microscopy Facility using a Titan Krios transmission electron microscope (Thermo Fisher Scientific) operated at an accelerating voltage of 300 kV. Movies were recorded on a K3 direct electron detector with a pixel size of 0.414 Å and a total accumulated electron dose of 50 eLJ/Å^2^. In total, 13,959 movies were acquired.

Cryo-EM data for the H4K12me1 nucleosome were collected at the Van Andel Institute Cryo-Electron Microscopy Facility using a Talos Arctica transmission electron microscope (Thermo Fisher Scientific) operated at an accelerating voltage of 200 kV. Movies were recorded on a K3 direct electron detector with a pixel size of 0.46 Å and a total accumulated electron dose of 50 e−/Å^2^. In total, 5,432 movies were acquired.

### Refinement and Model Building

All cryo-EM data processing was performed using cryoSPARC. For the complex between RomA and the H3K14Nle nucleosome, raw movie stacks were motion correction and binned by a factor of 2 (0.828Å/pix) followed by contrast transfer function (CTF) and motion correction. Particle picking was initially performed using blob picker and picked particles were extracted with an additional binning factor of 3 (2.484Å/pix). 2D classification was used to filter out junk particles and generate templates for template-based particle picking. Template-picked and blob-picked particles were re-extracted at bin 2 (1.656 Å/pix), merged, and filtered by 2D classification heterogeneous refinement. This refinement produced a single class containing a well-resolved nucleosome core. Further 3D classification without alignment yielded a single class with density on top of the nucleosome corresponding to RomA, which was refined to ∼4.2Å using the homogenous refinement job. To identify any stabilizing contacts with the nucleosome core, local refinement on the nucleosome core particle was conducted with a focus mask encompassing the nucleosome which refined to a final resolution of ∼3.8Å.

For the H2K12me1 nucleosome, raw movie stacks were motion correction and binned by a factor of 2 (0.94Å/pix) followed by contrast transfer function (CTF) and motion correction. Particle picking was initially performed using blob picker and picked particles were extracted with an additional binning factor of 3 (2.82Å/pix) Templates were generated for template-picking followed by particle extraction at bin 3 (2.82Å/pix). Multiple rounds of 2D classification and heterogeneous refinement of the template-picked particles resulted in a final map that refined to ∼6.0Å resolution. Particles from the final map were used to pick additional particles using Topaz^50^. Topaz-picked particles and template-picked particles were merged, re-extracted at bin 2 (1.88 Å/pix) and duplicates were removed. Multiple rounds of 2D and heterogeneous refinement yielded a final map at ∼5.2Å resolution.

