## Supplementary material for "Histone modification crosstalk between host and pathogen": Figure S

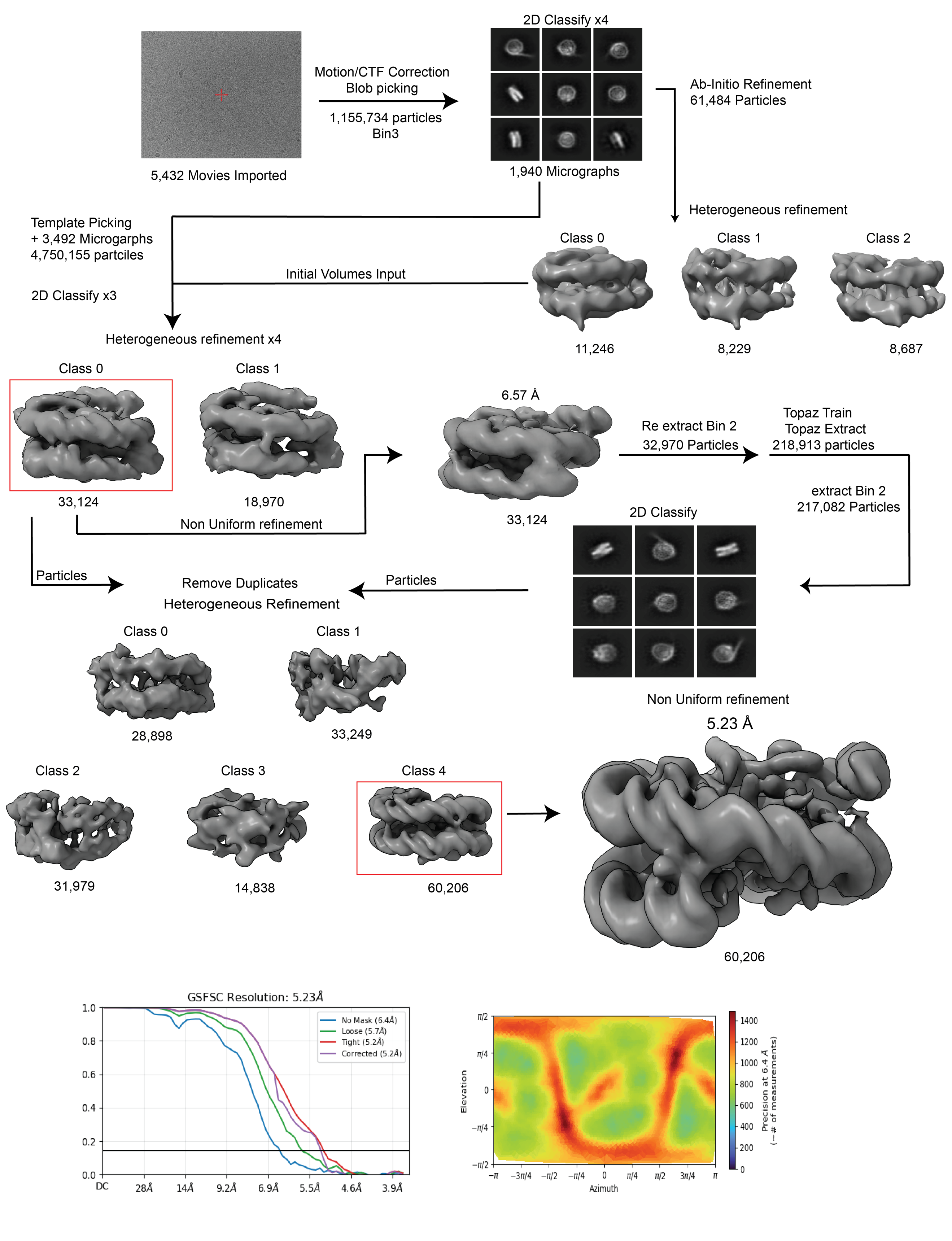

**Figure S1: Cryo-EM processing pipeline for the H4K12me1 nucleosome**

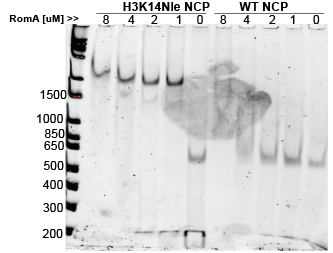

**Figure S2: H3K14Nle NCP purification and crosslinking**

7% Native Page showing RomA bound to Wildtype Nucleosome vs H3K14Nle Nucleosomes in the presence of SAM. RomA = 0-8uM, SAM = 200uM, Nucleosome = 200nM

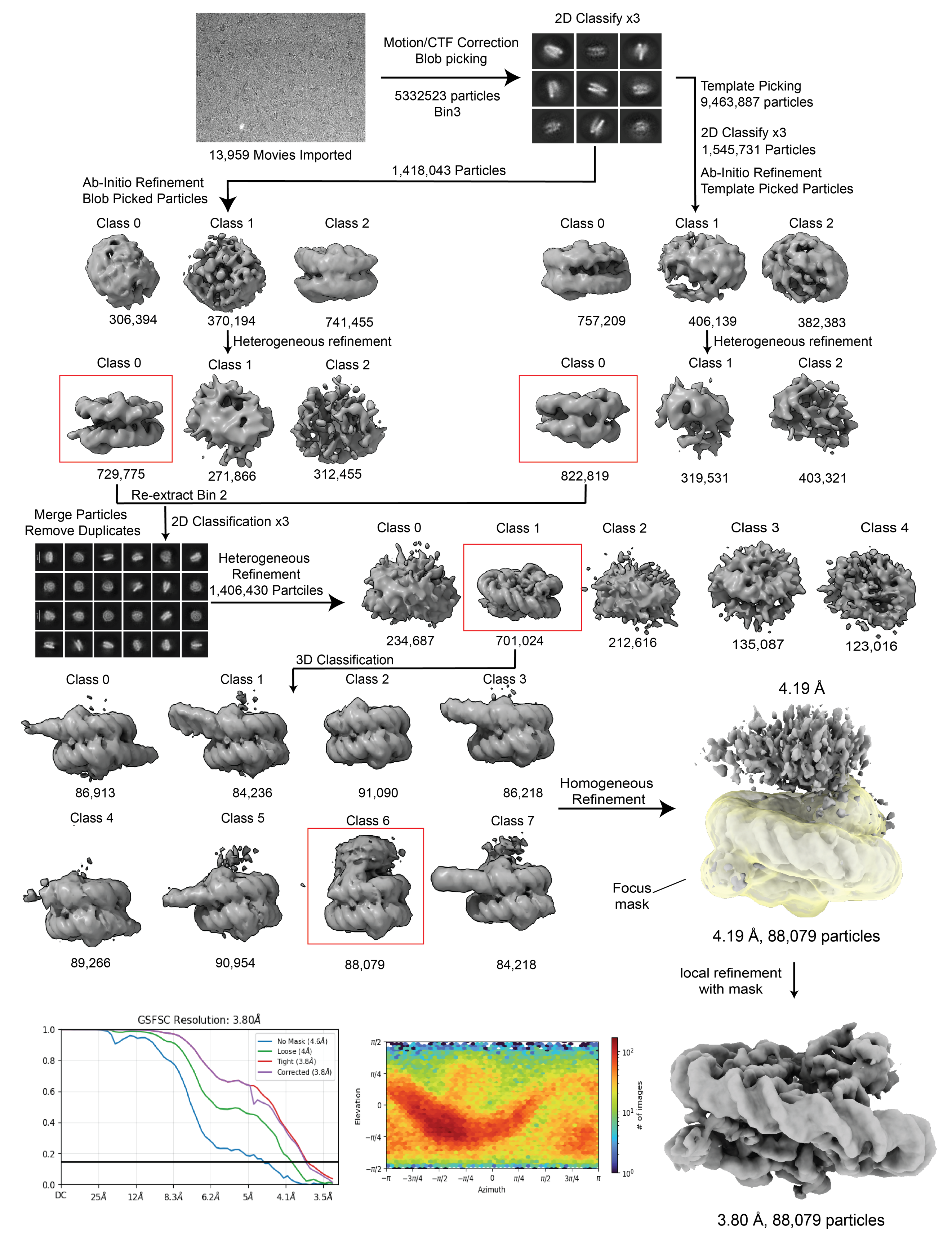

**Figure S3: Cryo-EM processing pipeline for the RomA H3K14Nle nucleosome complex**

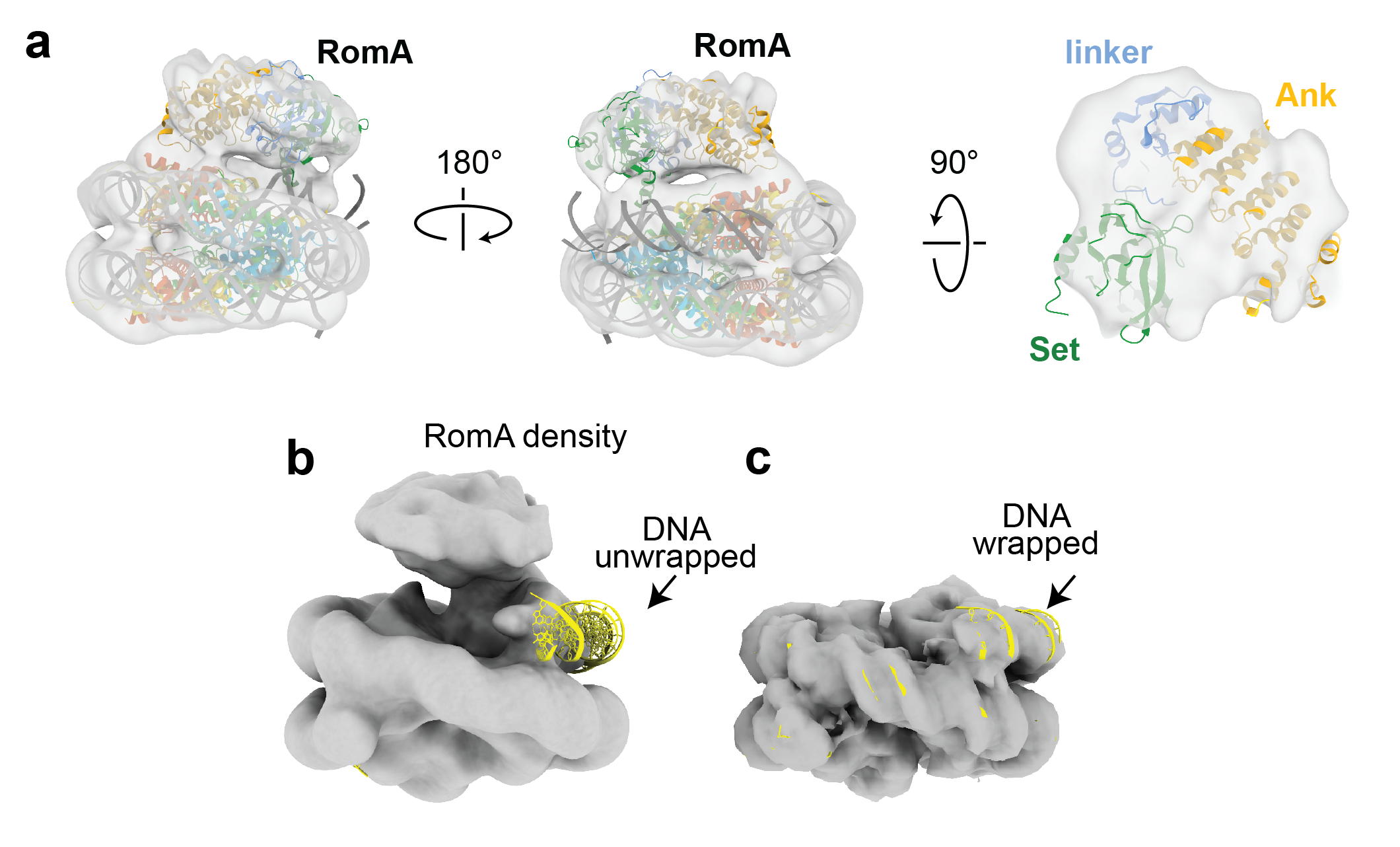

**Figure S4: Structural details of the RomA-nucleosome complex**

a) Cryo-EM structure of the RomA-([H3K14Nle]2) complex filtered to 15Å for clarity. The Cryo-EM density is shown as a semi-transparent gray surface and the structures of a nucleosome (PDB: 8YBJ) and LegAS4 (PDB: 5CZY) are docked into the Cryo-EM density. b) RomA-([H3K14Nle]2) complex filtered to 15Å showing that density for DNA is missing on the side of the nucleosome where RomA is observed. c) Cryo-EM density for the fully wrapped nucleosome (3D classification, Class 2, Fig. S3) where no RomA density is observed. DNA from the nucleosome (PDB: 8YBJ) is superimposed with the cryo-EM maps and colored yellow.

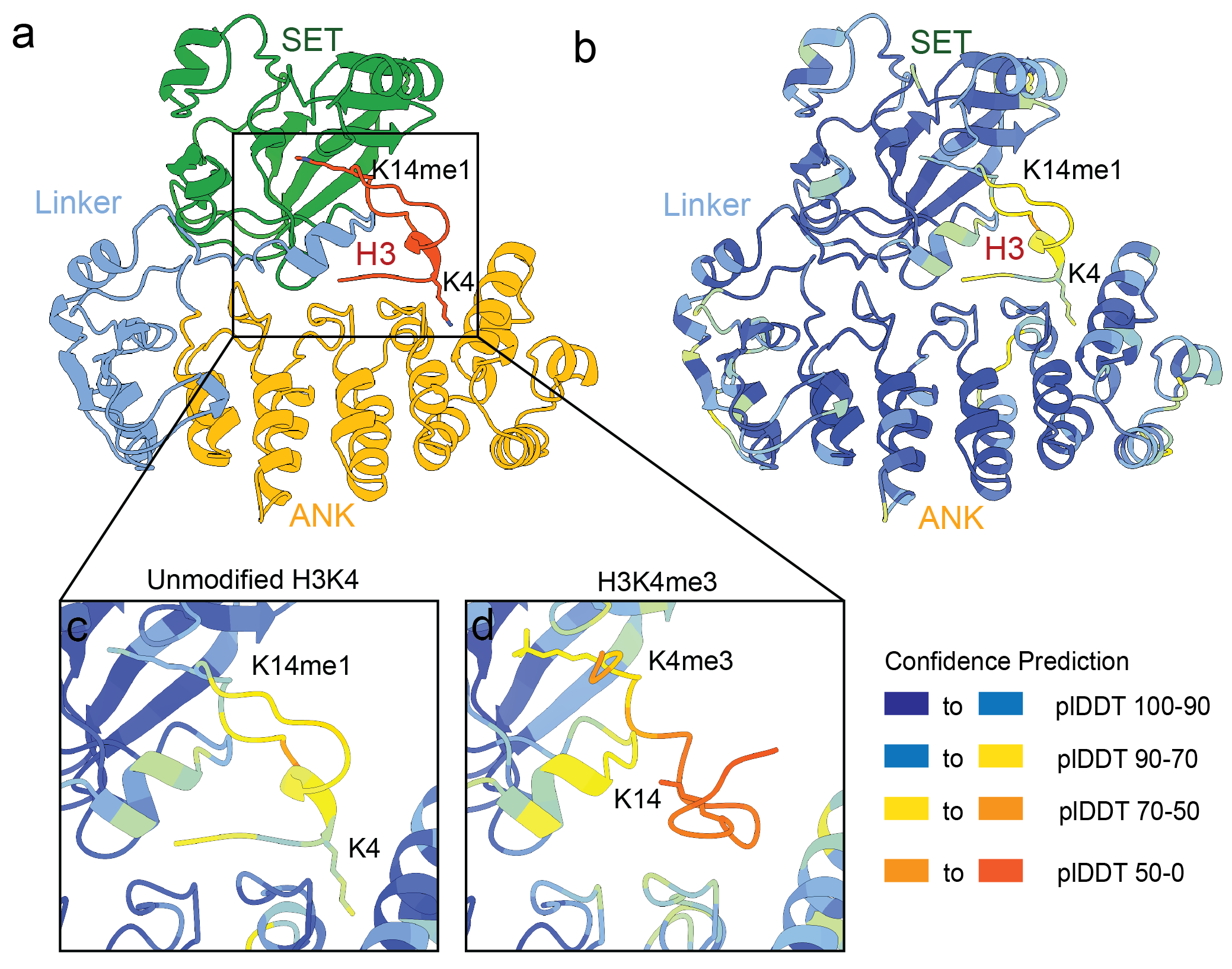

**Figure S5: AlphaFold prediction of RomA bound to an H3 peptides**

a) Alphafold prediction of RomA bound to a H3K14me1 (1-20) peptide colored by domains. H3K4 and H3K14me1 are depicted as sticks. b) Alphafold prediction as in a but colored by PlDDT scores. c) Close up view of the H3K14me1 peptide.
