## Supplementary material for "Histone modification crosstalk between host and pathogen": Table 1

| **Sample** | H4K12me1 nucleosome | RomA H3K14Nle complex: nucleosome focus | RomA H3K14Nle complex |
| --- | --- | --- | --- |
| **Data Statistics** |  |  |  |
| **Microscope** | Talos Artica | Titan Krios | |
| **Voltage (kV)** | 200 | 300 | |
| **Electron exposure (e^–^/Å^2^)** | 50 | 50 | |
| **Pixel size (Å)** | 0.46 | 0.414 | |
| **Symmetry imposed** | none | none | |
| **FSC threshold** | 0.143 | 0.143 | |
| **Map resolution range (Å)** | 25-5.2 | 25-3.8 | |
| **Initial particle projections** | 4750155 | 1406430 | |
| **Focus Region** | none | Nucleosome | none |
| **EMD code** | TBD | TBD | TBD |
| **Final particle projections** | 60206 | 88079 | |
| **Map resolution (Å)** | 5.2 | 3.8 | 4.2 |

**Table 1: Cryo-em data and refinement statistics**
